# Siccuracy: An R-package for executing genotype imputation strategy simulations with AlphaImpute

**DOI:** 10.1101/236760

**Authors:** Stefan McKinnon Edwards

## Abstract

**Background:** The reported R-package provides an easy way for executing and evaluating genotype imputation studies, by providing functions for preparing input files for AlphaImpute and efficiently calculating imputation accuracies. Using the correlation between true and imputed genotypes is used here as it is directly related to the accuracy of genomic prediction using imputed genotypes. This R-package calculates both correlation and counts correct and incorrect imputed genotypes.

**Results:** Implementing the correlation using a Fortran resulted in faster calculations and using less memory than using base R functions. Reporting the performance of an imputation should not be done only by the average correlation between true and imputed genotype. It is demonstrated that the highest average correlation is not necessarily the best correlation and that the range of obtained correlations provides a more nuanced grasp of the performance of the imputation.

**Conclusions:** An R-package is available that provides a fast, standardized, and tested implementation for computing the correlations.

## Background

The main motivation for this R-package was to provide a standardized and tested set of routines in R for genotype imputation with AlphaImpute (Hickey *et al*. 2011; Hickey, Kinghorn, *et al*. 2012; Antolín *et al*. 2017). These routines comprise preparing files for AlphaImpute and evaluating the resulting imputed genotypes, while avoiding loading extensive genotype data into memory. These routines therefore ease the task of implementing imputation strategy simulations that e.g. examines cost-benefit of different genotyping densities vs. accuracy of imputation.

This manuscript is structured as follows: First, the merits of genotype imputation is discussed and how it is typically evaluated. Code standardization is then briefly discussed, before introducing the AlphaImpute format which is related to some common genotype formats, followed by a more detailed description correlations as imputation accuracies. In Implementation, a one-pass-through method for calculating correlations is presented, followed by some details on our implementations. The results and discussion presents a simulation of an imputation scenario, discusses how to present imputation accuracies, before finally presenting the performance of our routines.

### Genotype imputation

Genotype imputation is a cost-effective method for obtaining large volumes of high density genotypes. In both genomic research and genomic prediction of livestock breeding values, individuals are routinely genotyped at different single-nucleotide-permutation (SNP) densities, e.g. the bovine 54K and 777K SNP chip (Matukumalli *et al*. 2009, 2011). This occurs as a subset of individuals are genotyped at high density (e.g. > 100,000 genetic SNPs) using expensive SNP chip arrays or genotype-by-sequencing (Gorjanc *et al*. 2015), while the remaining individuals are genotyped at lower densities (e.g. 500 – 50,000 genetic SNPs) at lower cost. The individuals genotyped at low densities are subsequently imputed to high density (Ma *et al*. 2013; Cleveland and Hickey 2013).

Imputation of missing genotypes has been implemented in a large range of software, such as PHASE (Stephens *et al*. 2001), fastPHASE (Scheet and Stephens 2006), Beagle (Browning and Browning 2007), SHAPE-IT (Delaneau *et al*. 2011), Impute2 (Howie *et al*. 2009), MaCH (Li *et al*. 2010), MERLIN (Abecasis *et al*. 2002), cnF2freq (Nettelblad *et al*. 2009), and AlphaImpute (Hickey *et al*. 2011; Hickey, Kinghorn, *et al*. 2012; Antolín *et al*. 2017). These rely on heuristic or probabilistic models to infer the missing genotypes by identifying the most probable haplotype (i.e. sequence of genotypes) that the individual has inherited from a close ancestor (heuristic model) or distance relative (probabilistic model) (Antolín *et al*. 2017). The softwares occupies different niches of the genomic research landscape, as some perform better when provided with pedigrees (uses heuristic models), as is common in livestock breeding, while others perform better when no pedigrees are available (Friedrich *et al*. 2017), as is common in human medicine. This paper will however not discuss the advantages and performances of the available imputation software packages, but refer to e.g. Ma et al. (2013) or Antolín et al. (2017) for this.

Evaluating the correctness of an imputation is typically done by counting the number of correctly and incorrectly imputed genotypes (Hickey, Crossa, *et al*. 2012; Calus *et al*. 2014), or by calculating the Pearson correlation between true and imputed genotypes (Calus *et al*. 2014). The latter is of interest in estimating livestock breeding values, as the expected accuracy of a breeding value estimated with imputed genotypes is directly proportional to this correlation (Mulder *et al*. 2012). Other measures exists such as error rate, yield (proportion of genotypes called), imputation quality score (Lin *et al*. 2010), or Hellinger score (Roshyara *et al*. 2014; Roshyara and Scholz 2015), alongside platform specific measures (e.g. those from MACH and IMPUTE), but these will not be covered here. Estimating the correctness of an imputation thus naturally requires knowing the true genotypes, so the correctness is estimated in simulation studies by *masking* known genotypes as missing.

Tested routines for masking datasets and estimating correctness are necessary for reproducible studies. This R-package has therefore been fitted with routines that allow researchers to efficiently run imputation strategy simulations by providing functions that are *human readable* and *tested*. Providing human readable functions eases a later review of the codebase, while testing ensures that the code does what is was designed to do. A more thorough case for this is presented in the next section. At the time of writing, no other R-packages have been found that offers these functions. Although R-packages for *imputing* genotypes do exist (e.g. ‘HIBAG’, https://github.com/zhengxwen/HIBAG; ‘alleHap’, https://cran.r-project.org/package=alleHap; ‘GeneImp’, https://pm2.phs.ed.ac.uk/geneimp/), this package differs by providing a framework around an existing genotype imputation software.

### Code standardization

The second aspect of the motivation for this R-package was to provide a fast, standardized, and tested implementation for computing the correlations. This minimizes the risk of sneaky bugs that otherwise can creep in when the source code of a presumed working implementation is duplicated to other projects, or by obfuscating the actual purpose of a code block. These issues are discussed in two letters by Wilson et al. (2014, 2016). An example of the latter is presented in the following code:

~~~
true <-as.matrix(read.table(‘TrueGenotypes.txt’, header=FALSE, row.names=1))
imputed <-as.matrix(read.table(‘ImputedGenotypes.txt’, header=FALSE,
row.names=1))
m <-apply(true, 2, mean)
s <-apply(true, 2, sd)
true <-scale(true, m, s)
imputed <-scale(imputed, m, s)
results <-sapply(1:nrow(true), function(i) cor(true[i,], imputed[i,]))
~~~

The above code assumes the individuals in the two input files have the same order, and that the two matrices are conformable. Determining the contents of the variable results can be tricky, especially considering the (deliberately) badly chosen variable name. The only hint to what results is lies in the middle of the anonymous function supplied to sapply. This code-example will calculate the correlation between rows of the two input files, and is a simplified version of the unit-testing for the function `imputation_accuracy`.

The approach in the above code example requires that both matrices are stored in memory in their entirety. Storing large datasets naïvely in R can be problematic (Rosario 2010; Kane *et al*. 2016). The approach in the first two lines is also sub-optimal as the function read.table itself can use large amounts of memory, and data duplication for converting the read data. frame to a `matrix` (Wickham 2014 chap. 17).

Lastly, additional code is required to handle missing values, mismatching rows, or datasets too large to store in memory. Each line of code added to handle special cases introduces potential bugs, for which bug fixes might not be forwarded to a subsequent project.

These issues are addressed by a) modularization, b) using meaningful names, c) implementation using a compiled language such as Fortran, and d) applying unit-testing to ensure correctness of implementation. The result can be called by a single function:

~~~
imputation_accuracy(‘TrueGenotypes.txt’, ‘ImputedGenotypes.txt’,
standardized=TRUE)
~~~

where the details of the function are discussed in detail later.

### AlphaImpute format and other, common formats

The file format for genotypes used by AlphaImpute is quite simple. It is a basic tabulated text file, space separated, where each line corresponds to an individual (i.e. sample) and consists first of an integer ID, followed by the *m* genotypes, either called (as integers 0, 1, or 2) or gene dosages (as numeric values between 0 and 2). Missing genotypes are usually coded with integer 9. A matrix stored in a simple text format is read as row-major, and is thus describe the AlphaImpute format as ‘ individual-major’. In contrast, the PLINK (Chang *et al*. 2015; Purcell and Chang 2016) format identifies an individual with two fields, a family ID and a within-family (sample) ID. Both the text and binary formats of PLINK are ‘SNP-major’, as a single SNP across all individuals are stored on a single line (text format) or data block (binary format). Our package offers a routine for converting the PLINK binary files to AlphaImpute format, with optional filtering output on chromosome, minimal allele frequency, SNP IDs, individuals, and automatic conversion of individual IDs; see `convert_plink`.

In addition, three other genotype formats are supported by the R-package: VCF (Danecek *et al*. 2011), Oxford (.gen and.sample files), and SHAPEIT (.haps and.sample file). These are also ‘SNP-major’ and all three text formats. The Oxford and SHAPEIT are very similar; Oxford records genotype probabilities for each state (two homozygote states and a heterozygote state), while SHAPIT records the presence of the alternate allele on each chromosome homologue.

The VCF format supports a very rich annotation of genotype data, but is not as much as standard as a promise of structured data. Support for this format is provided via the R-package ‘vcfR’ (Knaus and Grünwald 2016, 2017).

For the three latter formats, functions are provided for reading data into R, converting to the AlphaImpute format, and calculation of imputation accuracies.

### Imputation accuracy

There is currently no consensus for a definition of the ‘imputation accuracy’, so care must be taken when comparing ‘imputation accuracies’ between studies. ‘Imputation accuracy’ is here defined as the Pearson correlation between true and imputed genotypes, and can be applied to both called genotypes and gene dosages.

There are however three variations to this definition, the individual-wise, the SNP-wise, or the overall correlation: Given *n* individuals, each genotyped at *m* SNPs, define two matrices *A* and *B*, both of dimension *n* by *m*, where *A_ij_* describes the true genotype of the i^th^ individual at the j^th^ SNP, and *B* the same for the imputed genotype (either called genotype or gene dosage). The individual-wise (i.e. row-wise) correlation cor(*A*_*i*,1…*m*_, *B*_*i*,1…*m*_) describes how well the i^th^ individual has been imputed; the SNP-wise (i.e. column-wise) correlation cor(*A*_1…*n,j*_, *B*_1…*n,j*_) describes how well the j^th^ SNP has been imputed; and the matrix-wise correlation cor(*A, B*). Note that the three correlations result in an n-length vector, an m-length vector, and a scalar, respectively.

The expected genotype at different SNP positions vary due to differences in allele frequencies. Furthermore, rare alleles (SNPs where one allele is predominantly represented) are more difficult to impute correctly (Calus *et al*. 2014). *Standardization* of each SNP position by the allele frequency gives mores weight to SNPs with low allele frequencies; this standardization however only affects the individual-wise correlations, not the SNP-wise correlations.

Standardizing is performed separately on each column of matrices *A* and *B* using allele frequencies. In their absence, allele frequencies are estimated from the true genotypes in *A* and are applied to both matrices: The i^th^ column of *A* and *B* are thus standardized as 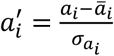 and 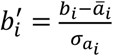, where *ā_i_* and *σ_a_i__*. are, respectively, the mean and standard deviation of the i^th^ column of *A*.

The individual-wise correlations are calculated for each individual across all SNPs of an individual. The Pearson correlation is invariant to scale and location for two random variables *X* and *Y*, i.e. for scalars *a, b, c*, and *d*, cor(*X, Y*) = cor (*a* + *bX, c* + *dY*), and it should be evident that 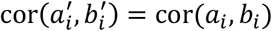. Standardizing these is performed with different means and standard deviations for each SNP, thus the above assumptions about the invariance to scale and location does not hold for the individual-wise correlations. It is later demonstrated that not standardizing the genotypes inflates the correlations. Calus et al. (2014) showed that without standardizing the genotypes, the imputation accuracies became highly biased for prediction accuracy of predicted breeding values.

In our implementation, standardizing is optional and with three options. The default method estimates the mean and standard deviation of each column of the ‘true’ dataset, and use this to centre and scale both datasets. As an alternative, the user may provide a vector of values to subtract from each column and a vector of values to divide each column by. Finally, a vector of allele frequencies, *p*, may be provided, in which case the centering and scaling are calculated as 2*p* and 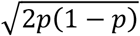, respectively. It is however important to note that the same values are used for standardising both matrices. If e.g. allele frequencies should be determined by a different set for individuals than in *A*, these can be estimated using the function heterozygosity (described later), with the added benefit of reuse of the estimates.

### Implementation

#### Pearson correlation

The Pearson correlation of two random variables, *X* and *Y*, can be written as three corrected sums of squares:

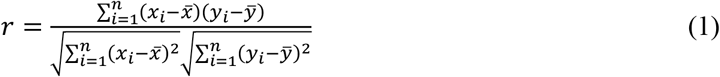

A naïve implementation of this is however not optimal as it requires 1) an initial run through each random variable to calculate the means (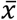 and 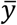), and 2) loss of precision due to round-off errors during the subtractions (Welford 1962). A one-pass-through algorithm was used, based on a recursive formula of the corrected sum of squares (Welford 1962), allowing the routine to calculate the three corrected sum of squares while only keeping the current data value (*x_i_* and *y_i_*) in memory in addition to a constant number of running statistics for each correct sum of squares. The algorithm is based on the following identities. Assuming a random variable *X*′ = (*x*_1_,…, *x_k_*), the corrected sum of squares is

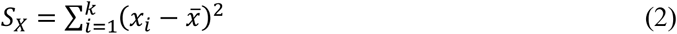

where 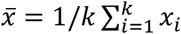. Welford (1962) showed that the corrected sum of squares of the first *n* ≤ *k* elements, 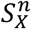, can be defined recursively as

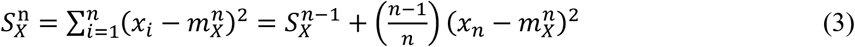

where 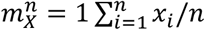 and 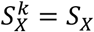. The running mean, 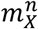, can in turn be defined recursively as:

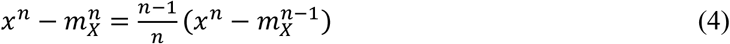

Expanding the right-hand square in eq. 3 and substituting with eq. 4, the corrected sum of squares for *X* can be written as

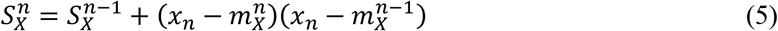

which is less susceptible to catastrophic cancellation and avoids a computational expensive division, although one is still required for calculating the mean. The statistics are initialised with 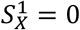 and 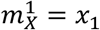.

The corrected sums of squares of both *X* and *Y* (nominator in eq. 1) can in turn be written as

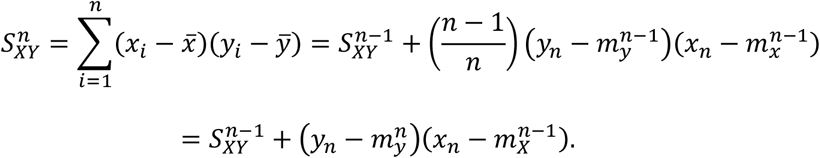

### Implementation of imputation accuracy

The imputation accuracy is implemented both for in-memory matrices in R and for files. The former was implemented to allow easy access to the same calculation for other formats, while the latter was implemented exclusively for the AlphaImpute format in two distinct Fortran subroutines: a non-adaptive subroutine with a low memory footprint, and an adaptive subroutine with a larger memory footprint. The adaptive subroutine has the advantage that the order of individuals in the two files do not have to be the same. In order to do so, it stores the entire dataset of the ‘true’ genotypes in memory. The non-adaptive subroutine only stores a single row of each file in memory as they are read. Both Fortran methods utilises the recursive formulas given above, and accepts the same arguments.

If using the default standardization, the entirety of the ‘true’ dataset needs to be processed prior to calculating the correlations. In the non-adaptive implementation, this causes the file of the ‘true’ dataset to be read twice, which incurs a running-time penalty from slower file I/O. The adaptive implementation will however estimate the column mean and standard deviation directly from the ‘true’ dataset kept in memory.

In R, the function `imputation_accuracy` was implemented as a generic method that allows method dispatching based on its first argument, which calls the file-based subroutines when passed a string (filename), the matrix-based method when passed a matrix, or even format-specific methods if passed an object obtained by reading e.g. a Oxford formatted file. The returned value is a list object of statistics per individual and per SNP, comprising correlation, scaling values, and number of correct, incorrect and missing genotypes in either true or imputed matrix, or both matrices.

### Heterozygosity

Heterozygosity refers to the frequency of individuals harbouring both alleles at a SNP position. If a population is homogenous, reproduces sexually, mates at random, and no selection nor mutation occurs, the allele frequencies can obtain Hardy-Weinberg equilibrium. Considering only a single SNP with a major allele frequency of *p*, under the Hardy-Weinberg equilibrium, the expected proportion of homozygote individuals for the major allele is *p*^2^, homozygote individuals for the minor allele is (1 – *p*)^2^, and heterozygotes 2*p*(1 – *p*). It is however departures from Hardy-Weinberg equilibrium that is usually of interest. The material given in this section is readily available on various online resources or textbooks, e.g. Lynch and Walsh (1998).

A simple allele-counting subroutine for diploids was implemented. For each column (i.e. SNP) of an input file, it counts the 0’s, 1’s, and 2’s (homozygote, heterozygote, and homozygote for other allele, respectively), while ignoring values outside this range. Values are read as integers, so if supplied with gene dosages (i.e. numeric values), the values are truncated to integers and thus might underestimate occurrences of 2’s. Denoting *n_i,het_* as the number of individuals heterozygotes for the i^th^ SNP, *n_i,hom_* the number of individuals homozygote for one allele, *n_i_* the number of non-missing genotypes, then the allele frequency for the i^th^ allele (*p_i_*), the observed heterozygosity (*H_i,obs_*) and expected heterozygosity (*H_exp_*) are calculated as

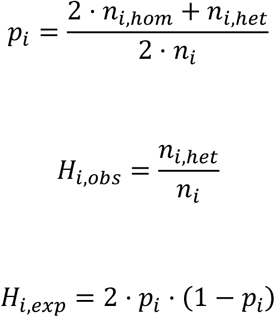

The observed and expected heterozygosity are routinely used in summary statistics, such as the inbreeding coefficient (*F* = [*H_exp_* – *H_obs_*]/*H_exp_*), or the fixation index (*F_st_*) which compares the heterozygosity between two subpopulations. For the latter, the implemented function accepts an argument that tallies the occurrences of heterozygotes and homozygotes separately for an arbitrary number of subpopulations. Note however that the summary statistics (*F* and *F_st_*) are given here without a subscript for SNP, as it is not trivial to extend these to multiple SNPs. See e.g. Bhatia et al. (2013) for a discussion on the subject.

The method is equivalent to that of PLINK (Chang *et al*. 2015; Purcell and Chang 2016) using the `--freq option`, but returns the result directly as a R data.frame without use of an intermediate file. PLINK currently only counts between two groups (case vs. control), whereas the function here allows for an arbitrary number of subpopulations.

### Converting PLINK binary files

`convert_plink` converts the PLINK binary bed/bim/fam files to AlphaImpute format, while allowing restriction on chromosomes, SNPs, minimal allele frequency, families, and individuals.

The PLINK binary format is very compact and stores genotypes as SNP-major, i.e. the first ⌈*n*/4⌉ bytes contains the first SNP, the next ⌈*n*/4⌉ bytes contains the second SNP, etc. A dataset of 4,342 genotyped dogs, genotyped at 160K SNPs (Hayward *et al*. 2016b, 2016a) takes up 166 MB on the disk. In the text format AlphaImpute uses, it is unpacked to 1.29 GB. In comparison to the PLINK binary format, the AlphaImpute is stored as individual-major. As the AlphaImpute format is individual-major, the entire dataset is stored in memory, before transposing and writing to file.

The conversion method has been implemented with some of the same functionalities as PLINK provides for restricting the output. This comprises filtering on minimal allele frequencies, chromosomes, SNPs, individuals and families.

Imputing genotypes is often performed on a per-chromosome basis. This, and the desire to avoid loading the entire data set into memory, prompted a preceding step in the conversion: splitting the PLINK binary file into multiple fragments, e.g. chromosomes, and then convert these fragments. This approach can be used by the argument `method=‘lowmem’ to convert_plink`.

The default behaviour of the `lowmem` approach is to split the dataset by chromosomes as given in the SNP map file with filename extension.bim. This can be modified with the `fragments` argument. When splitting the dataset into multiple `fragments`, the filenames of the resulting converted `fragments` can be given by the `fragmentfns` argument. The fragmentfns argument accepts either character vector for filenames or a filename generating function that accepts 0, 1, or 2 arguments. The first argument would be the fragment number (i.e. chromosome) and the second argument would be the maximum number of fragments. If there are not enough provided filenames, filenames of temporary files are generated with `tempfile(). tempfile()` does however not guarantee unique filenames if called by child processes of a forked R process.

### Testing correctness

All methods have been subjected to testing using the R-package testthat v. 1.0.2 (Wickham 2011). It was therefore easy to formulate anticipated scenarios in an easier readable programming language using base R functions and provide automated testing throughout development. For `imputation_accuracy` more than 1,200 lines of unit testing have been implemented, anticipating scenarios such as missing genotypes in one or both matrices, non-matching orders of individuals, or scaling parameters that include standard deviations of zero.

## Results & Discussion

### Simulating an imputation scenario

It is here demonstrated how to use the Siccuracy package for simulating an genotyping and imputation strategy, based on the public available dataset of Hayward et al. (Hayward *et al*. 2016b, 2016a). The dataset comprises 4,342 dogs of more than 150 breeds, mixed breeds, and free-ranging village dogs, from 32 countries worldwide. The dogs were genotyped at 185,805 SNPs, and the data is stored in a PLINK binary file set. 1,000 dogs are selected at random to be genotyped with a high-density chip array (HD), equal to the deposited data. The remaining dogs are simulated as being genotyped at a lower density (LD); 50, 300, and 1,000 SNPs per chromosome. For the demonstration, only the first chromosome (8,282 SNPs) is used. For each level of lower density, AlphaImpute v. 1.9 was used to impute the masked genotypes.

The simulation is as follows: Having downloaded the data and software, the genotypes are extracted, and for each pre-defined low-density, a subset of genotypes are masked for the low density genotyped dogs. After imputing the masked genotypes, the imputation accuracies can be calculated. The following displays a condensed block of code for the described simulation.

~~~
res <-convert_plink(‘cornell_canine’, outfn=‘cornell_canine_chr01.txt’)
method=‘lowmem’, extract_chr=‘1’, countminor = FALSE)
for (ld in low.densities) {
  mask.pos <-sample.int(hd, hd-ld, replace=FALSE)
  null <-mask_snp_file(fn=res$fnout, outfn=‘InputGeno.txt’, maskIDs=ld.dogs,
maskSNPs=mask.pos, ncol=hd)
  alphaimpute()
  result <-imputation_accuracy(res$fnout,
 ‘Results/ImputeGenotypeProbabilities.txt’, standardized=TRUE)
}
~~~

Data (Hayward *et al*. 2016b) was downloaded prior to the code as a PLINK binary file, and named ‘cornell_canine.bim’, ‘cornell_canine.bed’, and ‘cornell_canine.fam’. Chromosome 1 is extracted with `convert_plink. low.densities, ld.dogs`, and hd are vectors and a scalar whose defintions are hidden for brevity. `alphaimpute()` is a wrapper function for calling the external executables of AlphaImpute. Full setup with more verbose explanation can be viewed in Supplemental material S1.

Imputation accuracies were calculated both per animal and per SNP. The average imputation accuracy for the imputed dogs for the three low density chips were 0.18 (0.02 – 0.30), 0.48 (0.27 – 0.68), and 0.70 (0.57 – 0.84), with range between 1^st^ and 3^rd^ quantile given in parentheses. These are low imputation accuracies as neither selection of dogs or parameters for AlphaImpute were tuned for optimal imputation. AlphaImpute can in structured population achieve much higher accuracies, with values often up to 0.97 (Antolín *et al*. 2017). Figure 1 displays how the imputation changes for each imputed dogs as the number of SNPs of the low density is increased.

**Figure 1:**
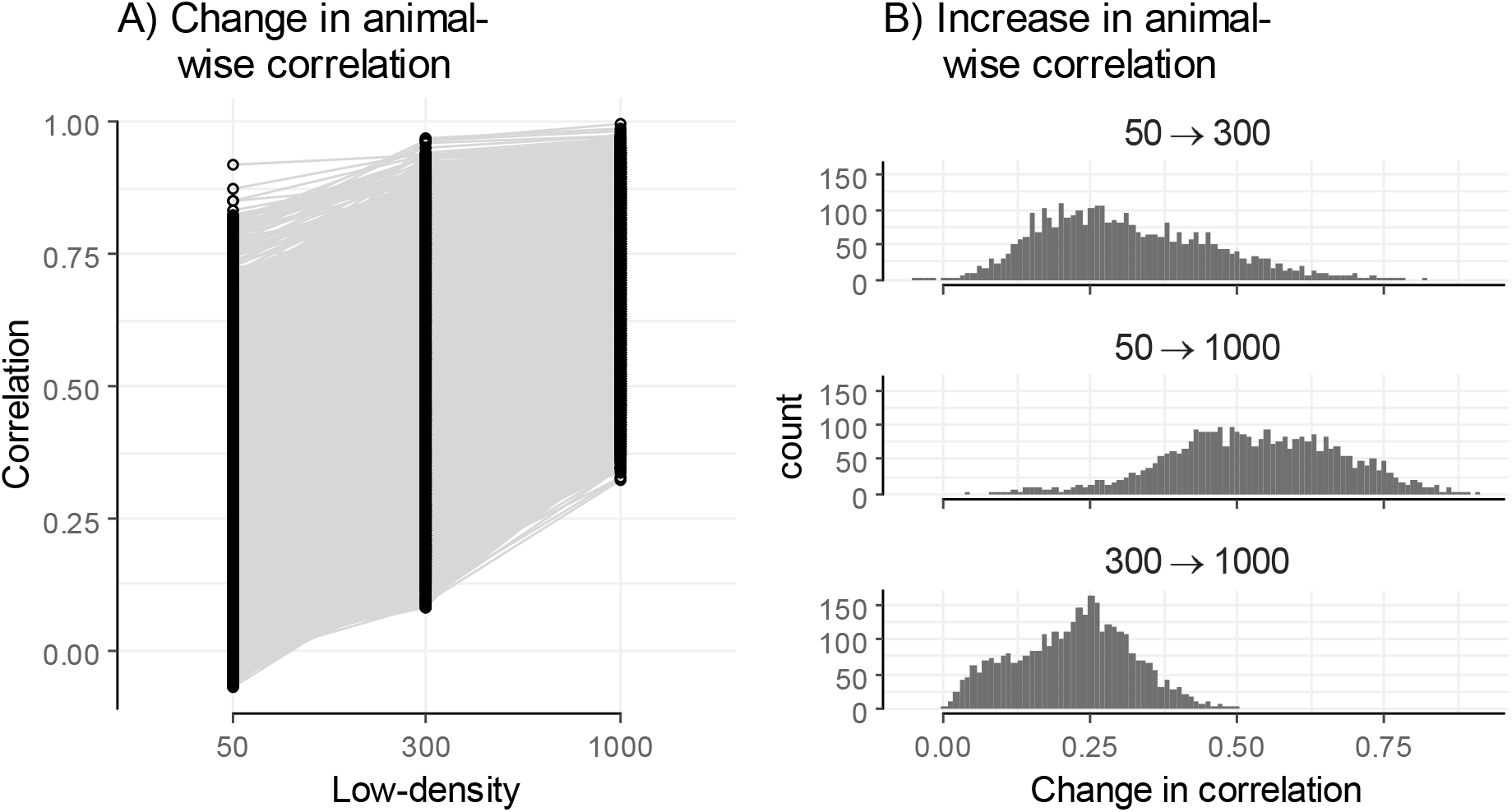
(Left) Animal wise imputation correlation at three different low-density genotyping options. Each dot represents an individual dogs and a grey line links the same dog, as imputed at three different low-densities. (Right) Histogram distribution of increase in imputation accuracy.

As noted above, standardizing the genotypes prior to calculating the correlations gives more weight to the alleles with lower allele frequencies. Figure 2 displays the trend of lower correlations with lower allele frequencies, concordant with Calus et al. (2014) that rare alleles are more difficult to impute. The tip at the lower allele frequencies could be due to population specific alleles, where close genetic similarities could allow correct imputation. The results displayed in Figure 2 are, as also noted above, invariant to the standardization, while the individual’s correlations are not, as displayed in Figure 3 where correlations are typically higher when standardization is not applied.

**Figure 2:**
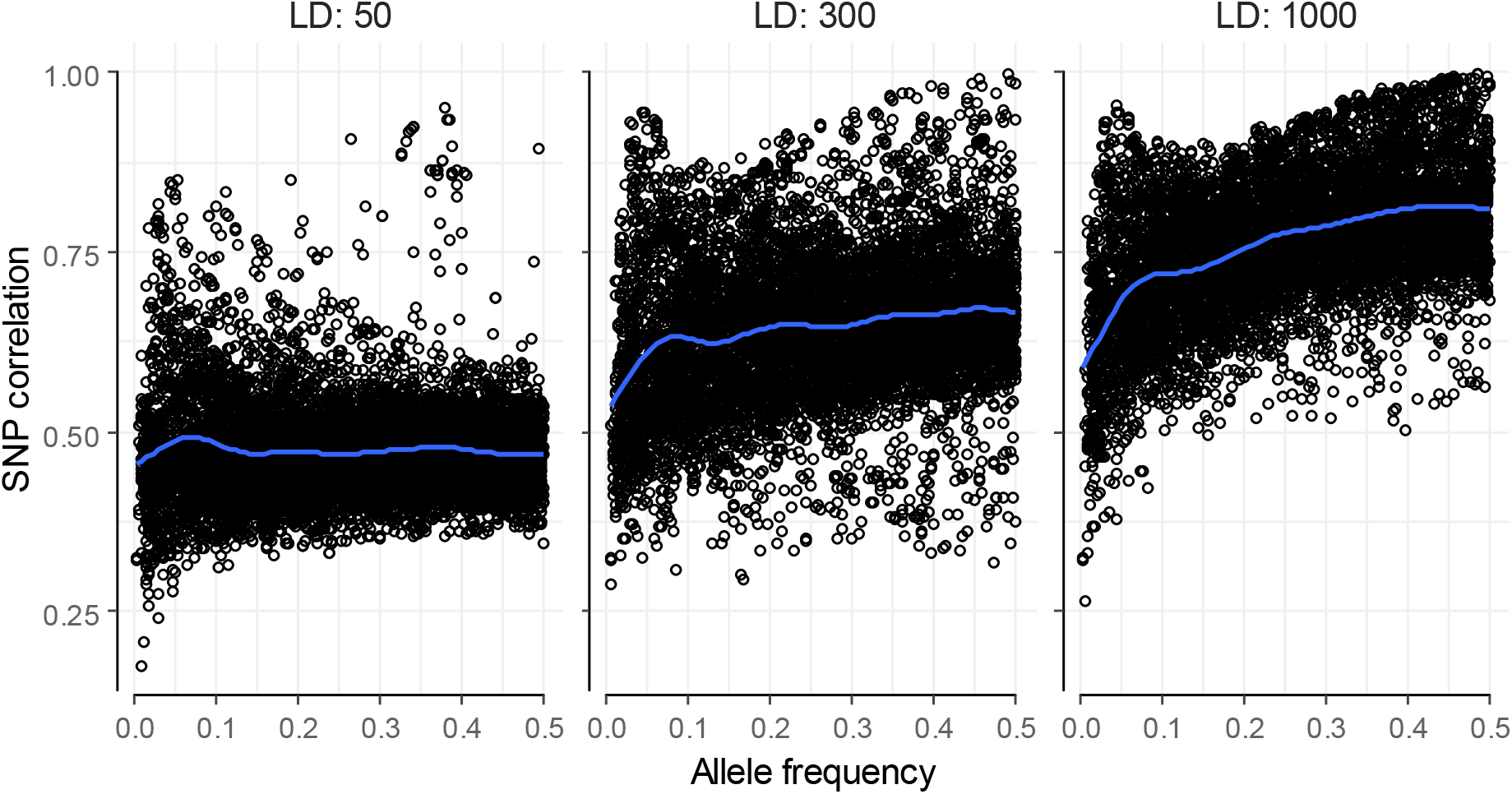
SNP-wise correlations of imputed SNPs as a function of allele frequencies, at different SNP densities. Blue line displays a smoothing spline.

**Figure 3:**
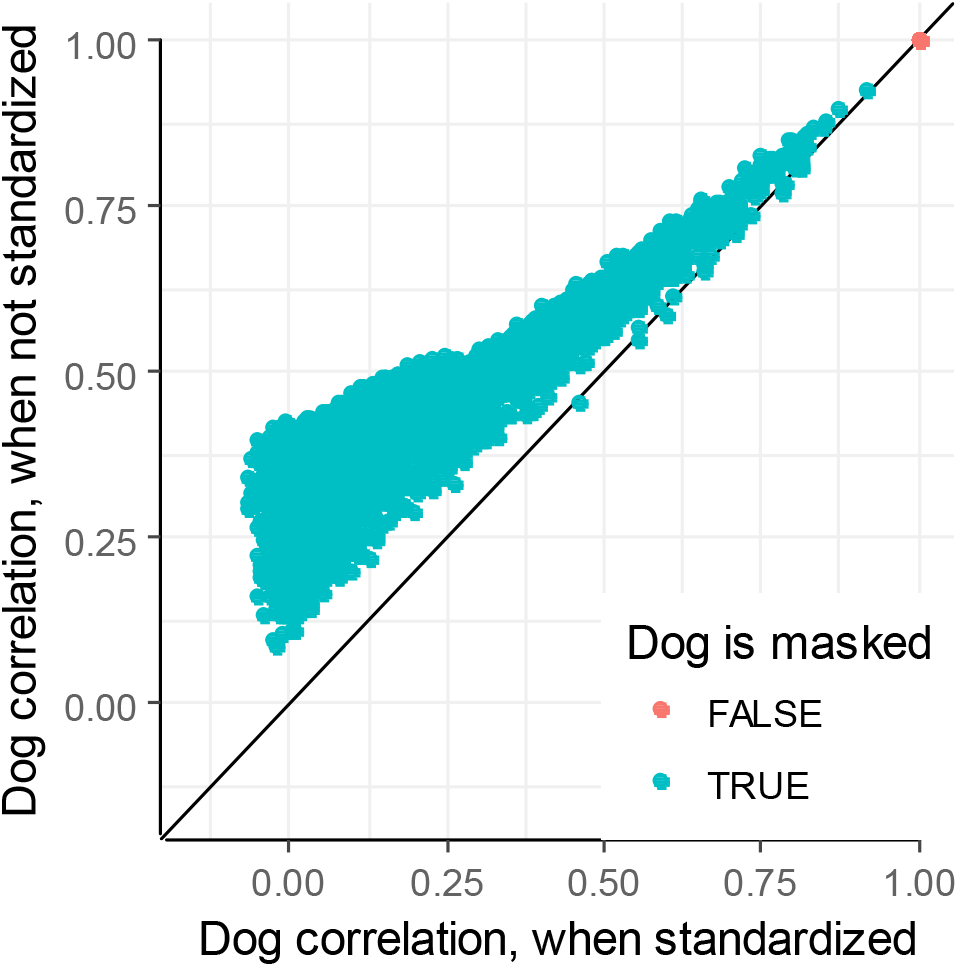
Correlations are inflated when genotypes are not standardized. Imputations were performed on dogs genotyped with 1,000 SNPs on chromosome 1 and subsequently imputed to 8,282 SNPs.

### Summarising correlations and the uncertainty of the estimated correlation

When reporting the imputation accuracies, it is often done by the average of all individual-wise or average of all SNP-wise correlations (Hickey, Crossa, *et al*. 2012; Ma *et al*. 2013; Calus *et al*. 2014). As these are realised values, the variance of these estimates may be of interest, especially for comparison between methods, but unfortunately often neglected (Erbe *et al*. 2012; Gorjanc *et al*. 2017).

The question remains how to best quantify the variance, uncertainty, or spread, of a correlation estimate. The uncertainty of a single correlation estimate, e.g. that for a single SNP, can be quantified by both theoretical approximations and bootstrapping (Cox 2008), which depend on the number of pairwise observations. Bootstrap estimates can be computational straining, while the theoretical approximations require a certain fidelity on the assumptions of bivariate normality. But what happens with the uncertainty of an average of correlations as those obtained for imputation accuracies?

Previous research (Cox 2008) has shown correlations (*r*) can be approximated by Fisher’s *z*-transformation *z* = atanh *r*, where *z* is normal distributed with mean atanh(*ρ*) + *c*(*ρ, n*) and variance 1/(*n* – 3) or 1/*n*. The correction *c*(*ρ, n*) depends on the approximation and is noted as 2*ρ*/(*n* – 1) or –5*ρ/2n*. With *n* in hundreds or thousands as in imputation accuracies, it can be neglected. Standard deviations calculated on z-transformed correlations are transformed back.

Standard deviations of randomly sampled correlations tend to asymptote 0, as the true correlation (*ρ*) approaches 1 (red line, Figure 4). Standard deviations of z-transformed correlations remain stable (blue line, Figure 4), as there is no limit on the z-transformed scale. The figure displays standard deviations of 1,000 realised correlations, produced by randomly drawing 1,000 pairs of observations from a standard bivariate normal distribution with known correlation (ρ).

**Figure 4:**
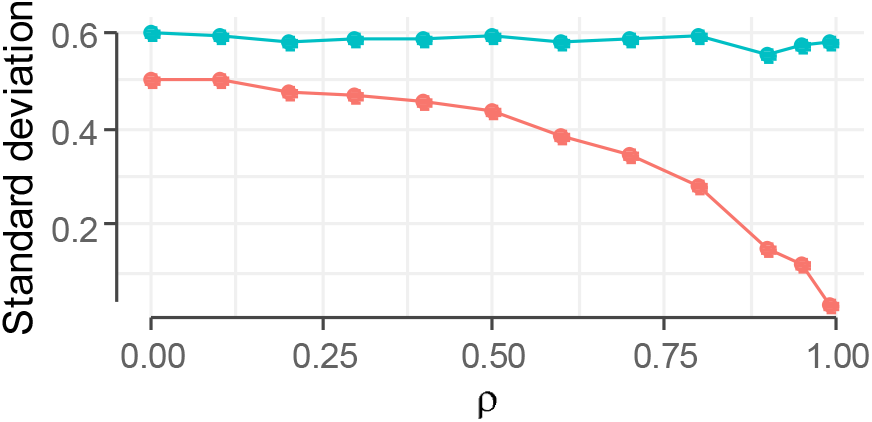
Standard deviations of z-transformed correlations (blue) are stable for all values of ρ, while those for untransformed correlations (red) asymptote towards 0 when ρ approaches 1. X-axis, ρ, is true correlation for standard bivariate normal distribution used for sampling correlations.

Both transformed and untransformed averages and standard deviations fail to capture the true correlations when *ρ* > 0.30 as displayed in Figure 5. For this, the 95% confidence intervals were calculated as 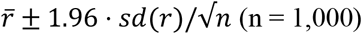. The displayed bias in Figure 5 are in line with those in (Corey *et al*. 1998) where averaging untransformed correlations underestimated true correlations, and averaging z-transformed correlations overestimated true correlations. Counter-intuitively, the bias *increases* with increasing number of correlations (Corey *et al*. 1998). As the average of the *untransformed* correlations are already widely used, this measure is used here in favour over the mean of the z-transformed correlations.

**Figure 5:**
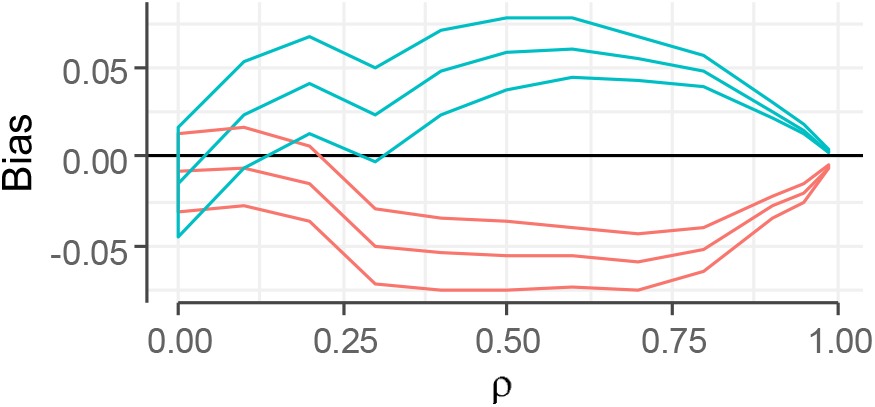
Bias of average estimate (average minus true correlation, ρ) with changing ρ when calculated on z-transformed (blue) or untransformed (red) correlations. For each ρ, 1,000 replicates of 1,000 pairwise observations were sampled. Lines, top to bottom, represent upper 95% confidence interval, mean, and lower 95% confidence interval.

These results were however obtained from independent samples from a nice behaving distribution. Imputation accuracies, whether calculated per individual across all SNPs or per SNP across all individuals, do not behave nicely, and depend on the linkage disequilibrium structure of the genome and relationship between individuals of those who are the basis for the imputation accuracies.

In this section forward, 3 sets of 2,000 individual-wise correlations from the imputation of the canine chromosome 1 in the previous section were used. The three sets, designated A, B, and C, were designed to have similar average, but different distributions. They are summarised in Table 1 and their histograms displayed in Figure 6.

**Table 1:** Mean, standard deviations (std.dev.), and percentiles of imputation accuracies in Figure 6.

|  | n | mean | std.dev. | 5% | 10% | 50% | 90% | 95% | mean† | std.dev† |
| --- | --- | --- | --- | --- | --- | --- | --- | --- | --- | --- |
| A | 2,000 | .745 | .142 | .490 | .560 | .758 | .937 | .965 | .497 | .8 |
| B | 2,000 | .776 | .102 | .609 | .633 | .784 | .909 | .929 | .290 | .8 |
| C | 2,000 | .758 | .125 | .541 | .580 | .772 | .912 | .952 | .433 | .8 |

†: Calculated on z-transformed correlations.

**Figure 6:**
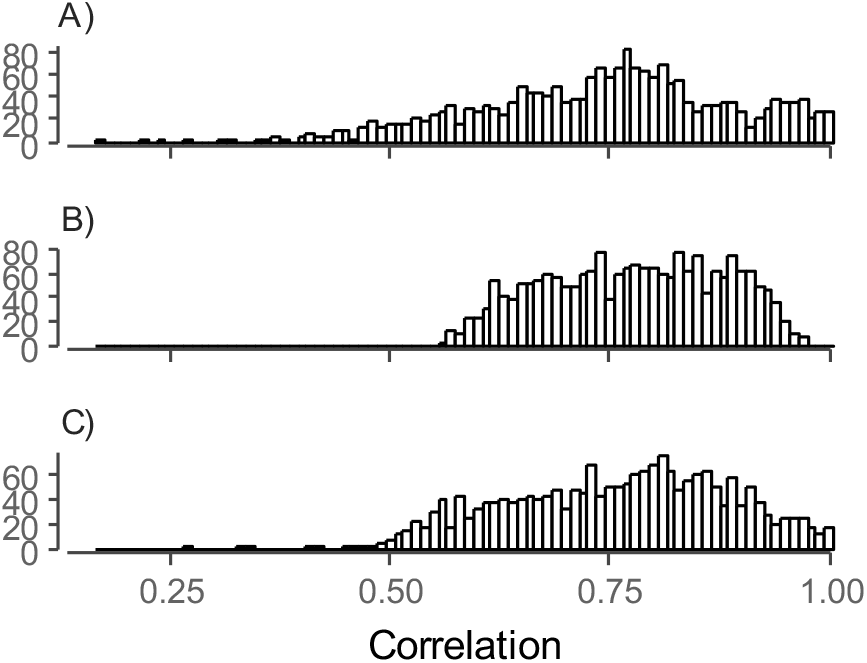
Histograms of three sets of imputation accuracies (n = 2,000) with similar mean and standard deviations.

When using the realised imputation accuracies, Fisher’s z-transformation may however inflate the standard deviation as 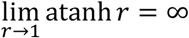. Figure 7 displays standard deviations of untransformed (red) or z-transformed (blue) correlations from set A in Figure 6. The correlations were binned in bins of size 0.05. The standard deviations of the untransformed correlations were as expected stable around the theoretical standard deviation of a uniform distribution due to being binned.

**Figure 7:**
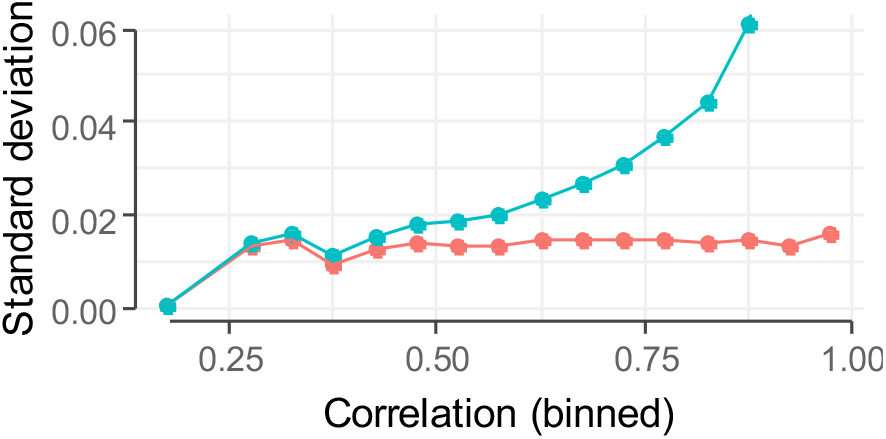
Standard deviations of z-transformed *SNP* correlations (blue) increases in bins of higher correlations (x-axis) whereas the standard deviation of untransformed *SNP* correlations (red) are stable, as expected. Realised imputation accuracies were binned in sizes of 0.05 and displayed on x-axis.

Figure 7 illustrates the issue of applying Fisher’s z-transformation on realised correlations from an unknown generative model / mixture of distributions, rather than sampled from a nice distribution. These results alone do however not discourage the use of the average and standard deviation for reporting. Above all, it is not advised to calculate the spread of correlations, when they are binned based on their value.

By measure of the mean *alone*, set B was the ‘best’ imputation as it has the highest mean. In addition, it has the smallest standard deviation. It is however evident from Figure 6 that it fails to produce imputation accuracies as high as set A and C. These have, respectively, 7.8% and 5.5% of imputation accuracies above 0.95, in contrast to 1.6% of set B.

The highest imputation accuracies were however obtained by set A, when evaluated on top percentiles (Table 1). This set however also has the longest lower tail, which explains its lower mean and larger standard deviation.

The order of the averages of z-transformed correlations is reversed (2^nd^ right column, Table 1) when compared to the average of the untransformed correlations. When calculating the average using z-transformed correlations, values closer to 1 are weighted heavier than those closer to 0. This is due to the non-linear transformation where 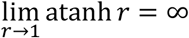, while being somewhat linear for *r* between −0.5 and 0.5. The average of the z-transformed correlations therefore better reflects the higher percentiles (90%, 95%, Table 1), where the average of the untransformed reflects the lower percentiles (5%, 10%, 50%, Table 1). Finally, the standard deviation of the z-transformed correlations betrays us in showing the three sets of imputation accuracies are contrived.

A long lower tail poses the question of why some SNPs or individuals are not imputed so well. An ungenotyped SNP may be difficult to impute if it lies far from a genotyped SNP, likewise an individual may be difficult to impute if it is the result of many meioses since the closest genotyped relative. The mean however does not reflect this proportion of less well imputed SNPs or individuals. An alternative to using an overall mean could be by summarising the individual imputation accuracies based on e.g. the individual’s traceability, such as in Mulder et al. (2012). An individual’s traceability quantifies the distance to nearest fully genotyped relative, based on a pedigree.

### Performance

It was assumed that calculations implemented in Fortran would be faster than using native R (R Core Team 2016). Both methods of correlations (adaptive and non-adaptive) were tested, by comparing performance of native R calculations, and calculations implemented in Fortran. The Fortran code was compiled in three different scenarios: 1) Compiled with the GNU Fortran (GCC) compiler when compiling R packages, ‘gfort’; 2) as 1, but with additional optimisations enabled with the –O3 flag, ‘gfort (O3)’; and 3) Compiled with the Intel IFORT compiler v. 16.0.0 with the –O3 flag, ‘ifort (O3)’.

Performance was estimated using the cpumemlog tools (Gorjanc 2015), which collects information via the linux ps command. Specifically, the statistics collected were ETIME (Elapsed Time) as measure of running time and RSS (Resident Set Size) as measure of memory usage. Random genotype files where generated with individuals *n* in range 100 – 20 000 and columns *m* in range 1000 – 30 000. The matrices were fully conformable and did not require re-ordering or matching of rows. 10 replicates of each size were used. Both correlations methods were evaluated, both with and without standardization. Performance was tested using R v. 3.3.3 on an array of Intel^®^ Xeon^®^ Processor E5-2630 v3 (2.4 GHz), as provided by Research Services at the University of Edinburgh.

The Intel IFORT compiled Fortran code was always quicker than GNU Fortran compile code, which in turn was always faster than native R (Figure 8). The figure depicts boxplots of the running time across 10 replicates of calculating correlations between matrices of 20,000 columns and 20,000 rows. There was no difference between the GNU Fortran compile code with or without optimisation. The Intel IFORT compiled Fortran code was faster by a factor of ~1.5 compared to the GNU Fortran code, while the latter was faster by factor of 1.5–2 compared to the native R. The slower running time of the GNU Fortran compile code of the non-adaptive method with standardization (top row, Figure 8) is likely due to the matrix being read in from disk twice, once to estimate standardization parameters and once to calculate correlations. The R code does not read in the file twice when standardizing, but still presents a longer running time than when not standardising.

**Figure 8:**
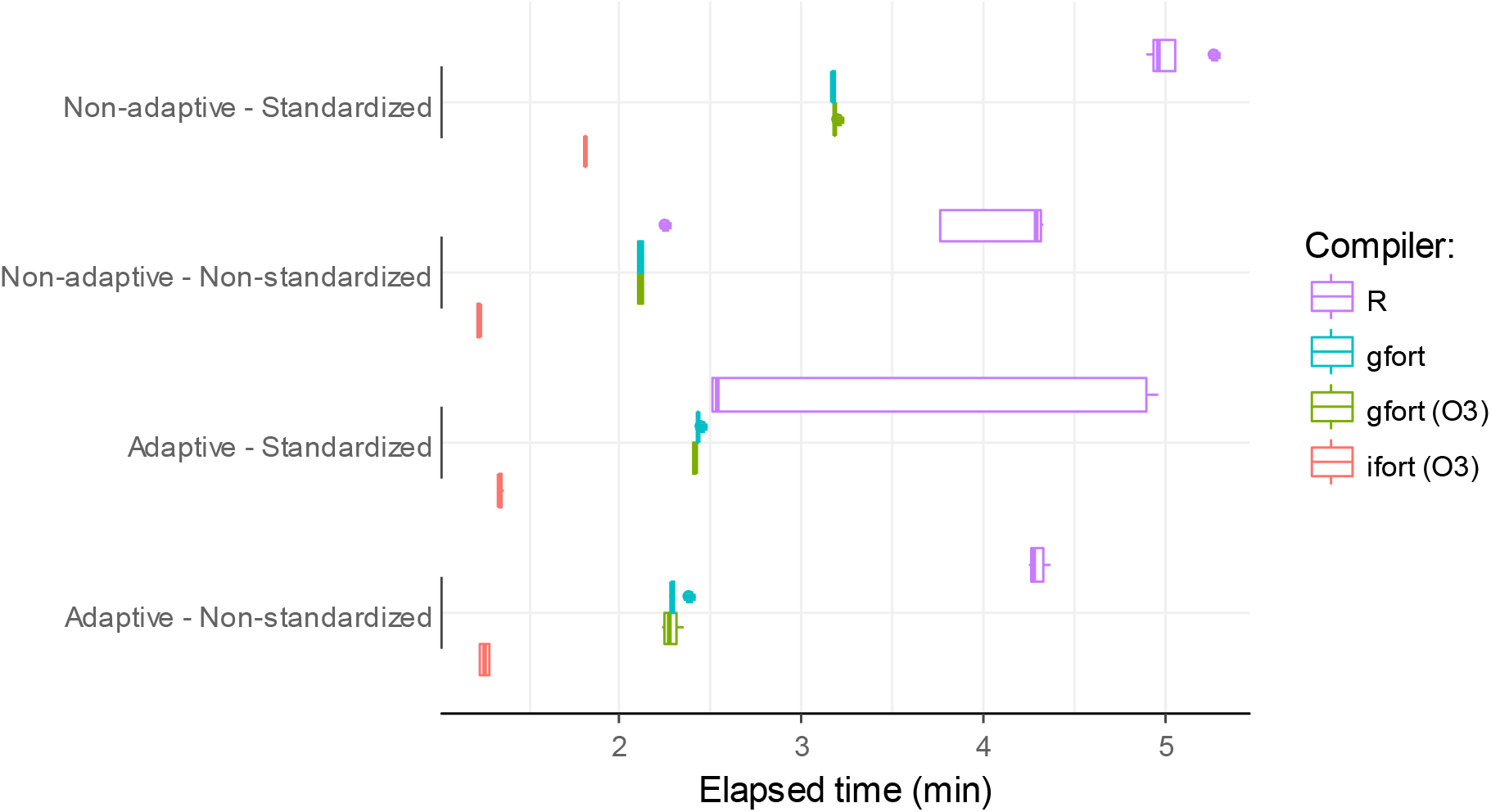
Elapsed time calculating correlations on 20,000 SNPs and 20,000 individuals.

Standardisation for the Fortran code required negligible time when the matrix is stored in memory, i.e. in the adaptive methods and in R (Figure 8). When not stored in memory, it almost doubles the running time. For all methods running time is as expected *O*(*nm*) (results not shown).

Calculating the correlations in R also takes a toll on the memory; for a dataset of 20,000 individuals and 30,000 SNPs, R required more than 12 GB of memory, as estimated by the RSS. In comparison, the Fortran libraries required about 32 MB for the non-adaptive method. If standardization is also required, R requires on average 14.7 GB memory, Fortran code libraries up to 34 MB. Based on this, the statistics for R are excluded from further comparisons on speed and memory usage.

The memory usage is *O*(*m*) for the non-adaptive method, and *O*(*nm*) for the adaptive method, where *m* is number of SNPs and *n* number of individuals (Figure 9). This is expected as the adaptive method stores the entirety of one matrix in memory, while the non-adaptive only stores a row. The top row of Figure 9 displays the low memory footprint of the non-adaptive method in magnitudes of MB. The high starting point (top-right panel, Figure 9) of approx. 25 MB is due to the measured memory consumption is of the entire R process, not just the subroutine.

**Figure 9:**
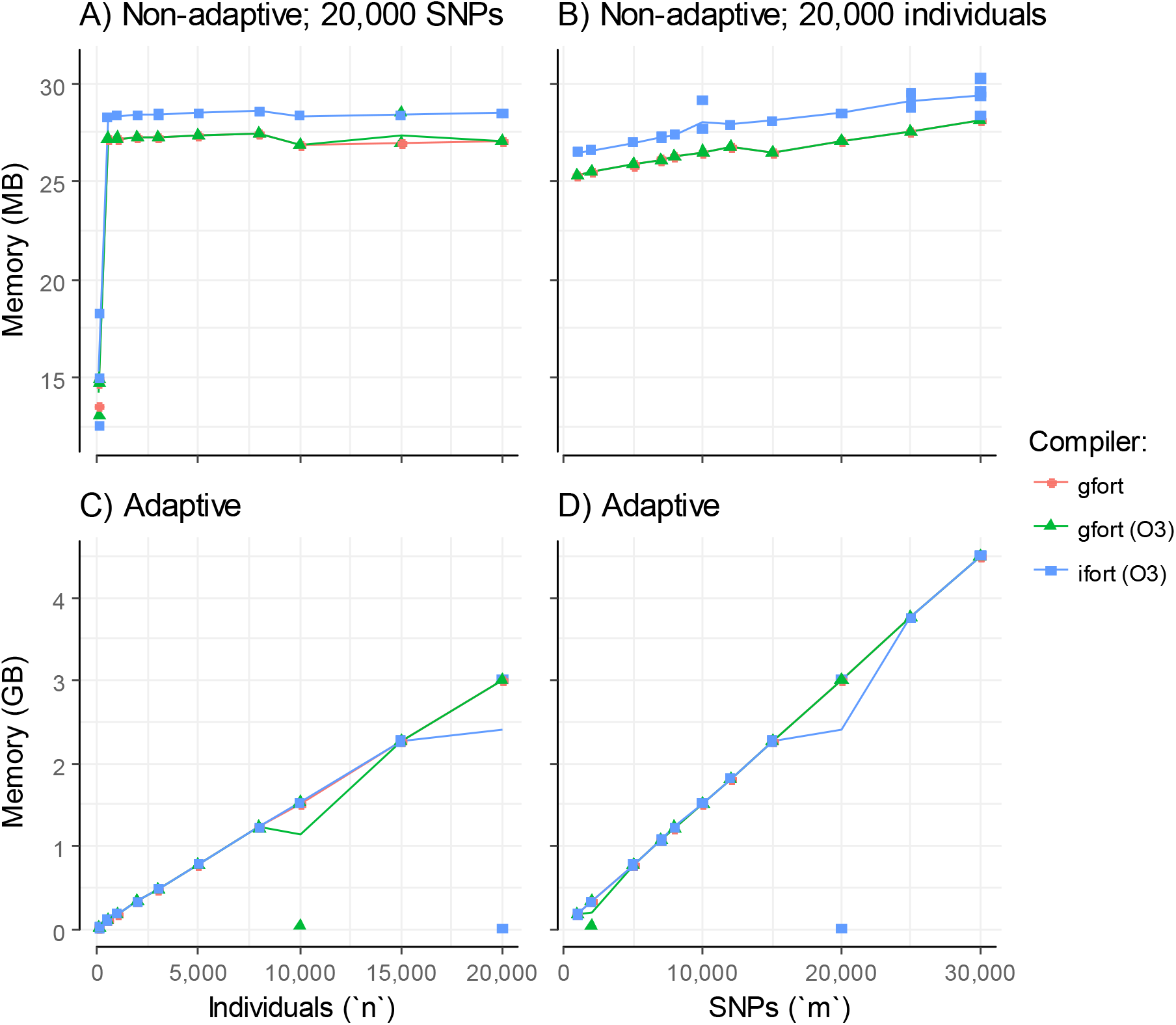
Memory usage of Fortran code as function of individuals (left column) or SNPs (right column). Top row is for the non-adaptive method with scale of memory in MB, bottom row is for adaptive method with scale of memory in GB. Data is for estimation without standardization.

There is little difference in memory usage of the different compilers, as seen by the overlapping colours in Figure 9. The Intel IFORT –O3 flag optimizes for speed and perform aggressive optimizations; the apparent cost of this is a slight increase in memory usage (blue line in top row, Figure 9).

The memory and time requirements of the calculations presented here are magnitudes smaller than those required for running AlphaImpute, whose task is vastly more complex. Preparing data and calculating correlations could therefore easily be performed on the same machine that performs the imputation, without having to consider restrictions on memory. This R-package has three distinct points of interest in this context: a) clearness of code, as presented in the beginning of this article, b) it allows preparation of the data on e.g. a local desktop computer, prior to submitting it to an imputation pipeline, and b) ad-hoc analysis, again on a local desktop computer.

## Conclusions

The Siccuracy R package presented in this paper has demonstrated increases in speed and reduction in memory usage by relying on Fortran coded subroutines. Further increases in speed were obtained by using the Intel IFORT compiler with optimisation. The R package has been designed to work directly with files, as AlphaImpute requires and in turn returns files.

Using the Pearson correlations as a measure of imputation accuracy is directly related to the reliability of estimated breeding values. Using the average and standard deviation of correlations may not necessarily be a statistic that describes the performance of the imputation well, as it does not reflect the tails of the distributions, where the lower tails may be of interest.

## Availability and requirements

Source code of the R-package is available on Github with instructions for installation at the project home page. A compiled zip-file for Windows users is also available.

Project name: Siccuracy

Project home page: https://github.com/stefanedwards/Siccuracy

Operating system(s): Platform independent

Programming language: R, Fortran

Other requirements: R (> v. 3.1.0)

License: GPL-3

Any restrictions to use by non-academics: Refer to License

## List of Abbreviations

LD: : Low-density (genotyping panel)
HD: : High-density (genotyping panel)
SNP: : Single nucleotide permutation

## Ethics approval and consent to participate

This study did not conduct new procedures for collecting genotype or phenotype information. For the dog genotype data used for demonstration, please refer to Hayward et al. (2016a) for full details. In short, blood samples were collected in accordance with the protocol approved by the Institutional Animal Care and Use Committee of Cornell University.

## Consent for publication

Not applicable

## Availability of data and material

The dataset analysed during the current study is available in the Dryad repository: http://dx.doi.org/10.5061/dryad.266k4 (Hayward *et al*. 2016b). Results generated during this study are available at the figshare respository: https://figshare.com/s/f48e9a85b6e03efca91f

## Acknowledgements

I wish to thank Daniel Money for valuable help in improving the written mathematical derivations of correlation.

## Supplemental material

### Supplemental material S1

canine_imputation.Rmd; knitr document demonstrating an imputation simulation study in dogs.

### Supplemental material S2

gfortran_vs_ifort.Rmd: knitr document for evaluating performance of correlation calculations.

